# An alpha 5-GABAa receptor positive allosteric modulator attenuates social and cognitive deficits without changing dopamine system hyperactivity in an animal model for autism

**DOI:** 10.1101/2023.08.24.554679

**Authors:** Adriana Jesus Souza, Dishary Sharmin, James M. Cook, Francisco S. Guimarães, Felipe V. Gomes

## Abstract

Autism Spectrum Disorders (ASD) are characterized by core behavioral symptoms in the domains of sociability, language/communication, and repetitive or stereotyped behaviors. Deficits in the prefrontal and hippocampal excitatory/inhibitory balance due to a functional loss of GABAergic interneurons are proposed to underlie these symptoms. Increasing the postsynaptic effects of GABA with compounds that selectively modulate GABAergic receptors could be a potential target for treating ASD symptoms. In addition, deficits in GABAergic interneurons have been linked to dopamine (DA) system dysregulation, and, despite conflicting evidence, abnormalities in the DA system activity may underly some ASD symptoms. Here, we investigated whether the positive allosteric modulator of α5-containing GABA_A_ receptors (α5-GABA_A_Rs) SH-053-2’F-R-CH3 (10 mg/kg) attenuates behavioral abnormalities in a rat model for autism based on *in utero* VPA exposure. We also evaluated if animals exposed to VPA *in utero* present changes in the ventral tegmental area (VTA) DA system activity using in vivo electrophysiology and if SH-053-2’F-R-CH3 could attenuate these changes. *In utero* VPA exposure caused male and female rats to present increased repetitive behavior (self-grooming) in early adolescence and deficits in social interaction in adulthood. Male, but not female VPA rats, also presented deficits in recognition memory as adults. SH-053-2’F-R-CH3 attenuated the impairments in sociability and cognitive function in male VPA-exposed rats without attenuating the decreased social interaction in females. Male and female adult VPA-exposed rats also showed an increased VTA DA neuron population activity, which was not changed by SH-053-2’F-R-CH3. Despite sex differences, our findings indicate α5-GABA_A_Rs positive allosteric modulators may effectively attenuate some core ASD symptoms.

## 1. Background

Autism Spectrum Disorders (ASD) are a group of neurodevelopmental disorders characterized by core behavioral symptoms in the domains of sociability, language/communication, and repetitive or stereotyped behaviors (Levitt and Campbell 2009, Fombonne 1999). The wide range of symptoms is proposed to result from various etiologies and, ASD are commonly considered multifactorial diseases, influenced by both genetic and environmental factors (Mandy and Lai 2016). Currently, no evidence-based pharmacological interventions are available for treating core ASD symptoms (Eissa et al. 2018). However, specific symptoms, such as agitation, irritability, and aggressive behavior, are often treated with the antipsychotics risperidone or aripiprazole, the only drugs approved by the FDA for children with ASD (Stepanova et al. 2017).

Epidemiological studies have indicated a positive correlation between prenatal exposure to valproic acid (VPA; also known as valproate) and the diagnosis of ASD (Weiss et al. 2009). Behavioral phenotypes related to core ASD symptoms, including impaired social behavior, repetitive or stereotyped behavior, and impaired communication, are modeled in rats and mice exposed to *in utero* VPA (Roullet, Lai and Foster 2013, Ornoy, Weinstein-Fudim and Ergaz 2016). The robust ASD-like phenotype induced by *in utero* VPA and its construct validity has made it a valuable rodent model for investigating the pathophysiology of ASD and exploring potential treatment targets (Vinten et al. 2009). Dysregulation of GABAergic neurotransmission in the brain has been implicated in several psychiatric disorders, including ASD (Bast, Pezze and McGarrity 2017). Deficits in the prefrontal and hippocampal excitatory/inhibitory balance due to a functional loss of GABAergic interneurons are proposed to underlie ASD symptoms (Helmeke et al. 2008, Zhang and Reynolds 2002, Grace and Gomes 2019, Bissonette et al. 2014, Nelson and Valakh 2015). These findings suggest that increasing the postsynaptic effects of GABA with compounds that selectively modulate GABAergic receptors could be a potential target for treating ASD symptoms (Cellot and Cherubini 2014).

Low doses of benzodiazepines, such as clonazepam, mitigate impairments in social interaction and cognitive function in animal models for autism (Han et al. 2012). However, the specific subtypes of GABA_A_ receptors (GABA_A_R) involved in these effects have not been elucidated. Targeting GABA_A_R, particularly by potentiating α2, α3, and α5 subunit-containing GABA_A_R, may produce therapeutic effects without benzodiazepine-related side effects (Prevot et al. 2019). Unlike α1-4 subunit-containing GABA_A_R, which are distributed throughout the brain, α5 subunit-containing GABA_A_Rs (α5-GABA_A_Rs) are primarily expressed in the hippocampus and, to a lesser extent, in the prefrontal cortex (Fritschy and Panzanelli 2014). Within these brain regions, α5-GABA_A_Rs are predominantly found on pyramidal neurons, where their activation modulates their firing rate, thus contributing to the regulation of the excitatory-inhibitory balance (Semyanov et al. 2004). Therefore, the potentiation of the α5-GABA_A_R-mediated neurotransmission may be effective in disorders where the excitatory-inhibitory balance is dysregulated, such as ASD.

Here, we investigated whether the α5-GABA_A_Rs positive allosteric modulator SH-053-2’F-R-CH3 attenuates behavioral abnormalities in the rat model for autism based on *in utero* VPA exposure. In addition, given that deficits in GABAergic interneurons have been linked to dopamine (DA) system dysregulation in rat models for mental illnesses, such as schizophrenia and Alzheimer’s disease (Eassa et al. 2023, Grace and Gomes 2019), and, despite conflicting evidence, abnormalities in the DA system activity are proposed to underly some ASD symptoms (Roullet et al. 2013, Dichter, Damiano and Allen 2012, Pavăl 2017, Chevallier et al. 2012), we also evaluate if animals exposed to VPA *in utero* present changes in the ventral tegmental area (VTA) DA system activity through in vivo electrophysiology and if SH-053-2’F-R-CH3 could attenuate these changes. Male and female rats were used to evaluate potential sex differences.

## 2. Material and methods

### Animals

Male and female Sprague-Dawley rats were obtained from the colony maintained by the Central Animal House of the University of São Paulo, campus Ribeirao Preto. Animals (1 male and 1 female) were housed together to copulate in micro-isolated cages, with water and food available ad libitum and under standard laboratory conditions. The male rat was removed after detecting spermatozoids in the vaginal smear, making it gestational day 0 (GD0). Pregnant dams were injected with saline or VPA (600 mg/kg) on GD12. Male and female pups were weaned on postnatal day (PND) 21±1 and housed two to three per cage. All procedures were approved by the Ribeirao Preto Medical School Ethics Committee, which follows Brazilian and International regulations.

### Drugs

VPA (Sigma, USA) was dissolved in saline and SH-053-2’F-R-CH3 (a positive allosteric modulator of α5-GABA_A_Rs synthesized by Dr. James Cook’s group, University of Wisconsin Milwaukee, USA) was dissolved in 2% Tween 80 in saline. All drugs were intraperitoneally (i.p.) injected in a 1 mL/kg final volume.

### Experimental Design

We first evaluated repetitive behaviors (spontaneous grooming) during early adolescence and adulthood in male and female rats exposed to VPA *in utero*. In adulthood, sociability and cognitive function deficits evaluated through the social interaction (SI) and novel object recognition (NOR) tests, respectively, and changes in VTA DA neuron activity were assessed. In a second set of experiments, we evaluated the effects of the positive allosteric modulator of α5-GABA_A_Rs SH-053-2’F-R-CH3 (10 mg/kg) in attenuating the behavioral and electrophysiological changes caused by *in utero* VPA in adult animals. SH-053-2’F-R-CH3 was administered 30 min before each behavioral test or electrophysiology recording. In the NOR test, SH-053-2’F-R-CH3 was administered 30 min before the retention trial. The dose of SH-053-2’F-R-CH3 was based on previous work showing that this compound attenuated behavioral changes in a rat model for schizophrenia based on neurodevelopmental disruption (Gill et al. 2011).

### Behavioral tests

#### Repetitive behavior (spontaneous grooming)

The time animals spent in grooming behavior was evaluated during early adolescence (at PND 30 for males and PND32 for females) and adulthood (at PND 62 for males and PND 64 for females). For this, animals were placed in a circular arena (60 cm diameter and 65 cm height) for 15 min. The test was videotaped, and the grooming time was measured manually. Locomotor activity was also evaluated using Any-Maze software (Stoelting, USA).

#### Social interaction (SI) test

Social interaction was evaluated in adulthood (at PND 62 for males and PND 64 for females). An experimental rat and an unfamiliar conspecific rat with the same sex and age were placed in the opposite sides of a circular arena (60 cm diameter and 65 cm height) that they could freely explore for 15 min.

The time of active social behaviors engaged by the experimental rat, such as sniffing, following, grooming, and climbing on or under the other rat, was recorded.

#### Novel object recognition (NOR) test

After the SI test, animals were assessed in the NOR test (at PND 63 for males and PND65 for females). First, animals were placed in a circular arena (60 cm diameter and 65 cm height) for 15 min for habituation. Twenty-four hours later, each animal was evaluated in two trials: acquisition (T1) and retention trial (T2). In the acquisition trial, each rat was placed in the arena containing two identical objects for 10 min. One hour later, animals were put back in the arena for 5 min for the retention trial (T2). For this trial, one of the objects presented in T1 was replaced by a novel, unfamiliar object. The behavior was recorded on video for blind scoring of object exploration. The time the animal spent directing its face to the object, up to 2 cm away from it, while watching, licking, sniffing, or touching it with the forepaws, was measured. Recognition memory was assessed using the discrimination index, corresponding to the difference between the time exploring the novel and the familiar object, corrected for the total time exploring both objects (discrimination index = [novel − familiar/ novel + familiar]).

#### *In vivo* recordings of VTA DA neurons

After the behavioral tests, animals (PND 66-80) were anesthetized with chloral hydrate (400 mg/kg, i.p.) and mounted on a stereotaxic frame. Animals were supplemented periodically with chloral hydrate to maintain suppression of the hindlimb withdrawal reflex. Body temperature was maintained at 37°C with a temperature-controlled heating pad. In vivo single-cell extracellular recordings of VTA DA neurons were conducted using single glass microelectrodes (impedance 6–8 MΩ) filled with a 2% Chicago Sky Blue solution in 2 M NaCl. Electrodes were placed in the VTA (anteroposterior: −5.3 mm from the bregma; mediolateral: +0.6 mm lateral to the midline, and −6.5 to −9.0 mm ventral from the brain surface) using a hydraulic positioner. DA neurons were identified according to well-established electrophysiological features (Ungless and Grace 2012). Three parameters were measured: population activity determined by counting the number of spontaneously firing DA neurons encountered while making 6–9 vertical passes within the VTA in a predetermined pattern, average firing rate, and the percentage of action potentials occurring in bursts (burst activity). Each identified DA neuron was recorded for 1-3 min. At the end of the recording, the electrode sites were marked via iontophoretic ejection of Chicago Sky Blue dye from the electrode (20 μA constant negative current, 15-20 min) for posterior histological confirmation.

### Statistical analysis

Data were presented as the mean ± SEM and analyzed using two or three-way ANOVA followed by Tukey’s post-test. p<0.05 was considered significant.

## 3. Results

### In utero VPA increases repetitive behavior during early adolescence and causes deficits in sociability and cognitive function in adult male and female rats

During early adolescence, male and female VPA rats presented increased self-grooming behavior (Figure 1A). Two-way ANOVA revealed a significant effect of VPA exposure (F_1,32_=23.90, p<0.05) and sex (F_1,32_=6.94, p<0.05), without interaction between them. Spontaneous grooming increased in male and female rats exposed to VPA (p<0.05 vs. saline; Tukey’s post-test). These changes did not persist until adulthood (Figure 1B). In addition, adolescent and adult VPA rats did not present differences in locomotor activity (Supplementary Figure 1).

**Figure 1.**
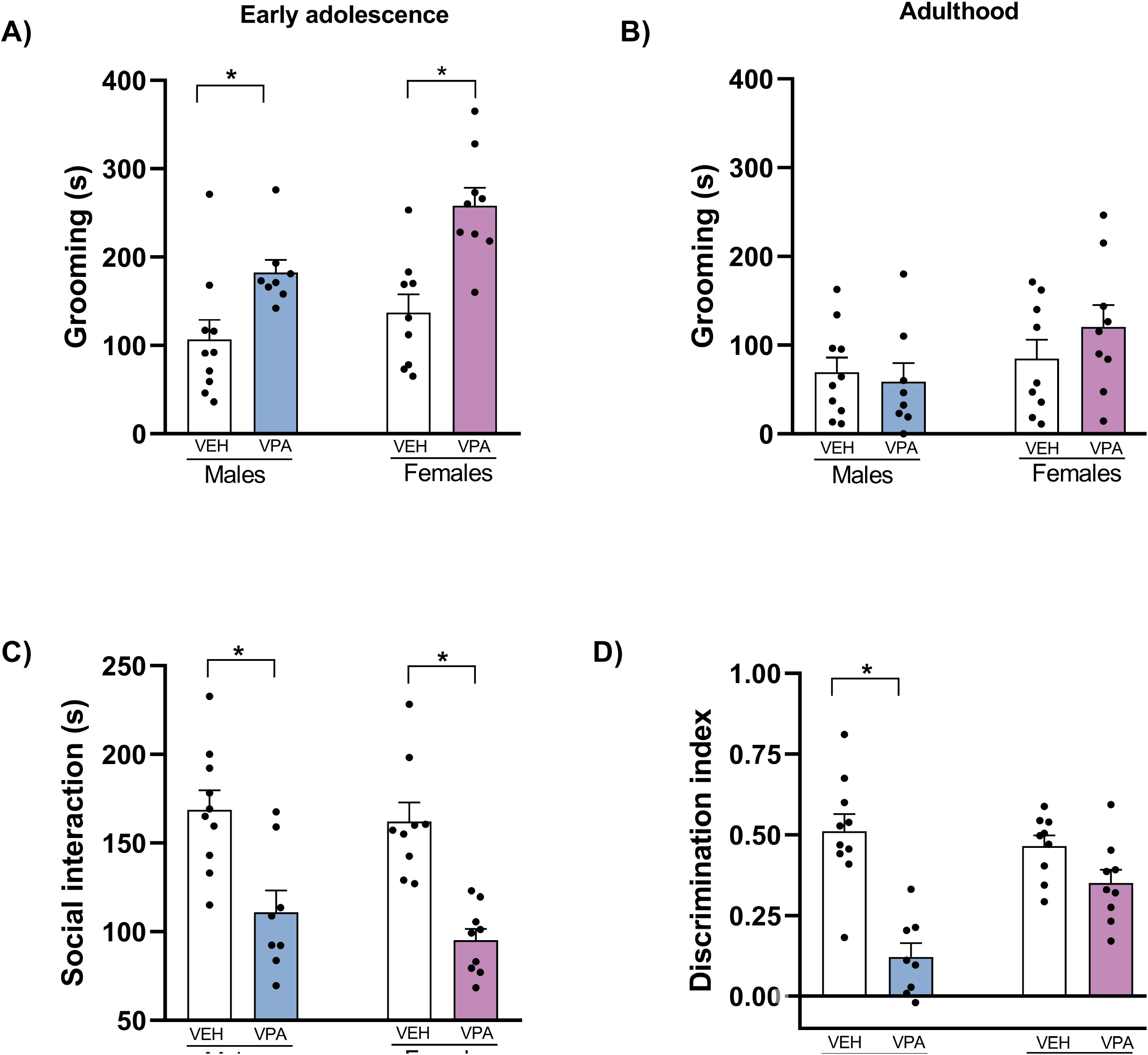
**(A)** *In utero* VPA exposure increased self-grooming time in males and females during early adolescence, **(B)** but not in adulthood. However, **(C)** a decrease in social interaction was observed in adult male and female VPA rats. Regarding cognition, **(D)** only male VPA-exposed rats presented a decrease in recognition memory in the NOR test (n=8-10/group). *p<0.05 vs. saline rats, two-way ANOVA followed by Tukey’s post-test.

Regarding sociability, adult male and female VPA rats showed decreased social interaction. A two-way ANOVA revealed a significant effect of VPA exposure (F_1,32_=36.25, p<0.05; Figure 1C) without effect of sex and interaction. Regardless of sex, VPA-exposed rats spent less time engaged in social interaction with an unfamiliar rat (p<0.05 vs. saline; Tukey’s post-test).

In the NOR test, there was no difference between the exploration of the identical objects placed on the right or left side of the arena during the acquisition session for all groups (Supplementary Figure 2A), indicating a lack of spatial preference. In the retention session, a greater exploration of the novel object was observed for all groups, except male rats exposed to VPA *in utero* (Supplementary Figure 2B). These findings were reflected in the discrimination index (Figure 1D). A two-way ANOVA showed a significant effect of VPA exposure (F_1,32_=32.52, p<0.05), sex (F_1,32_=4.27, p<0.05), and interaction between them (F_1,32_=9.66, p<0.05). Post hoc analysis indicated a decrease in the discrimination index in males exposed to VPA (p<0.05 vs. saline; Tukey’s post-test), but not in females. These findings indicated that deficits in the novel object recognition memory caused by *in utero* VPA exposure appears to be sex-depend since only males were affected.

### In utero VPA increases VTA DA system activity in adult male and female rats

One day after the behavioral experiments, we started the recordings of VTA DA neurons. An increased VTA DA neuron population activity was found in VPA rats, regardless of sex (Figure 2A). A two-way ANOVA indicated a significant effect of VPA exposure (F_1,20_=26.04, p<0.05), without effect of sex and interaction. Male and female VPA-exposed rats showed an increased number of spontaneously active VTA DA neurons (p<0.05 vs. saline). No change was observed for the firing rate (Figure 2B) and burst activity of the identified DA neurons (Figure 2C).

**Figure 2.**
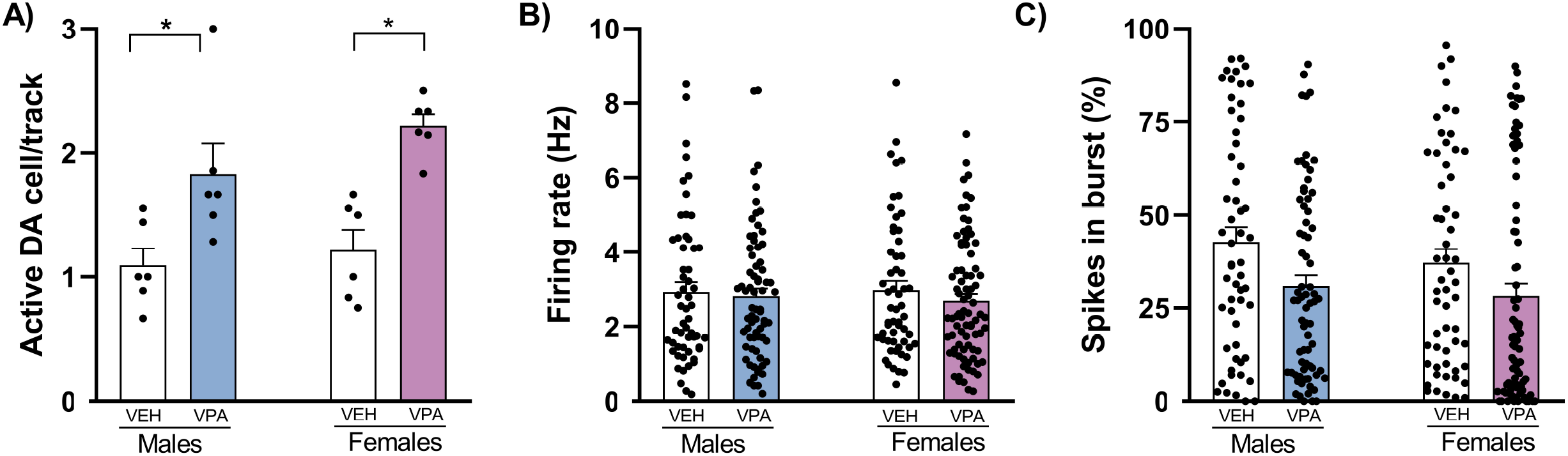
*In utero* VPA caused males and females to show **(A)** an increased number of spontaneously active VTA DA neurons (n= 6 animals/group), **(B)** with no change in the firing rate **(C)** and % of spikes in burst (males – saline: 56 DA neurons, VPA: 63 DA neurons; females – saline: 57 DA neurons, VPA: 82 DA neurons). *p<0.05 vs. saline rats, two-way ANOVA followed by Tukey’s post-test.

### SH-053-2’F-R-CH3 attenuates behavioral deficits in VPA rats without changing the increased VTA DA system activity

We tested the effects of SH-053-2’F-R-CH3 in attenuating changes in the social interaction and NOR tests and VTA DA system activity in adult male and female VPA rats. A three-way ANOVA indicated an interaction between sex and treatment in the social interaction (F_1,63_=4.59, p<0.05) and an interaction among sex, treatment, and VPA exposure in the NOR test (F_1,63_=5.69, p<0.05), suggesting effects mediated by sex in these tests. For VTA recordings no interaction with sex was observed. We further analyzed these data using 2-way ANOVA.

For the social interaction in male rats, a two-way ANOVA indicated an interaction between VPA exposure and SH-053-2’F-R-CH3 treatment (F_1,22_=4.32, p<0.05) and a trend for a significant effect of condition (F_1,22_=3.87, p=0.06). Tukey’s post-test demonstrated a significant decrease in social interaction behavior in the vehicle-treated VPA rats (p<0.05 vs. vehicle-treated saline), which was not shown by SH-053-2’F-R-CH3-treated VPA rats (p>0.05 vs. vehicle-treated saline; Figure 3A). In females, a two-way ANOVA revealed a significant effect of VPA exposure (F_1,32_=7.10, p<0.05), without an effect of treatment and interaction. Tukey’s post-test indicated a significant decrease in social interaction in female vehicle- and SH-053-2’F-R-CH3-treated VPA rats (p<0.05 vs. vehicle-treated saline; Figure 3B). These findings indicate that SH-053-2’F-R-CH3 attenuated sociability deficits in male VPA rats but not females.

**Figure 3.**
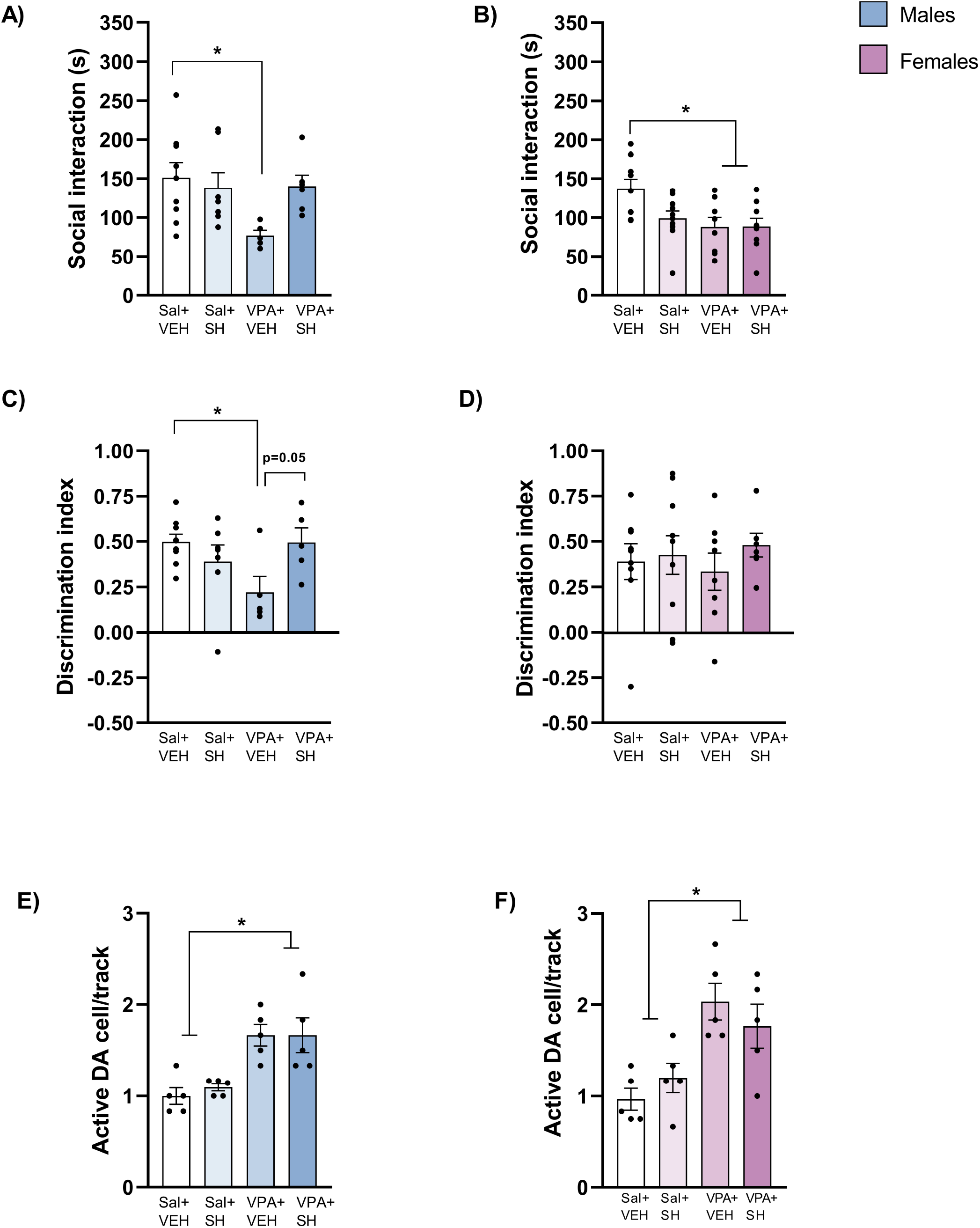
**(A)** SH-053-2’F-R-CH3 attenuated impairment in the social interaction test in male, **(B)** but not in female rats exposed to VPA. **(C)** SH-053-2’F-R-CH3 also reverted deficits in novel object recognition memory in male VPA rats. **(D)** Female VPA rats did not present deficits in the NOR test. In addition, the increased VTA DA neuron population activity found in **(E)** male and **(F)** female VPA rats was not changed by SH-053-2’F-R-CH3 (behavioral tests -n=5-10 animals/group; VTA recordings -n=5/group). *p<0.05, two-way ANOVA followed by Tukey’s post-test.

In the NOR test, there was no difference between the exploration of the identical objects during the acquisition session for all groups (Supplementary Figure 3A). In the retention session, a greater exploration of the novel object was observed for all groups, except male vehicle-treated VPA rats (Supplementary Figure 3B). The discrimination index reflected these findings (Figures 3C and D). In males, a two-way indicated a significant interaction between SH-053-2’F-R-CH3 treatment and VPA exposure (F_1,22_=9.45, p<0.05), without treatment and interaction effects. Post hoc analysis showed a decrease in the discrimination index in vehicle-treated VPA rats (p<0.05 vs. vehicle-treated saline; Tukey’s post-test), with a trend to be reversed by the treatment with SH-053-2’F-R-CH3 (p=0.059 vs. vehicle-treated VPA; Tukey’s post-test). As we previously found, female VPA rats did not present deficits in the discrimination index (Figure 3D).

For the VTA DA system activity, SH-053-2’F-R-CH3 did not attenuate the increased VTA DA neuron population activity in male and female VPA rats (Figures 3E and F). Two-way ANOVA indicated a significant effect of VPA exposure in males (F_1,16_=25.61, p<0.05) and females (F_1,16_=19.38, p<0.05), without treatment and interaction effects. Tukey’s post-test indicated, independent of the treatment and sex, VPA rats showed a significant increase in the number of spontaneously active VTA DA neurons (p<0.05 vs. vehicle-treated saline; Tukey’s post-test). No significant changes were observed for the firing rate and burst activity (Supplementary Figure 4).

## 4. Discussion

*In utero* VPA exposure caused male and female rats to present increased repetitive behavior (self-grooming) in early adolescence and deficits in social interaction in adulthood. Male, but not female rats, exposed to in utero VPA also presented deficits in recognition memory as adults. The treatment with the α5-GABA_A_R positive allosteric modulator SH-053-2’F-R-CH3 attenuated the impairments in sociability and cognitive function in male VPA rats, but without attenuating the decreased social interaction in female VPA rats. In addition to the behavioral abnormalities, male and female adult VPA rats showed an increased VTA DA neuron population activity. Regardless of sex, SH-053-2’F-R-CH3 did not alter the enhanced VTA DA system activity caused by *in utero* VPA.

Repetitive behavior, including self-grooming, has been described in the VPA autism model in different developmental periods (Kang and Kim 2015, Mehta, Gandal and Siegel 2011, Qi et al. 2022), but few studies have investigated the same animal longitudinally (Fereshetyan et al. 2021). Here, we evaluated the same animal during early adolescence and adulthood and found increased grooming time only during adolescence. The causes for the absence of changes in adult VPA animals in our study are unclear. However, it may involve habituation since grooming was evaluated in the same apparatus in the two periods. It was found that adult mice exposed to in utero VPA do not show repetitive self-grooming in a familiar environment (Kang and Kim 2015).

Similar to previous studies (Dai et al. 2018, Chaliha et al. 2020), we also found that adult VPA rats presented sociability and recognition memory impairments. Notably, novel object recognition memory deficits were not observed in female VPA rats, indicating sex differences in the performance of VPA rats in the NOR test. These findings agree with reports indicating that VPA-exposed female rats show better performance in some cognitive tasks (Ghahremani et al. 2022, Melancia et al. 2018).

Abnormalities in GABAergic neurotransmission have been implicated in ASD (Port et al. 2017, Zhang et al. 2020). Evidence from different animal models for autism, including the VPA model, has suggested that functional deficits of GABAergic interneurons are linked to behavioral abnormalities associated with autism, such as stereotyped behaviors and deficits in social interaction and cognitive function (Lauber, Filice and Schwaller 2016, Fontes-Dutra et al. 2018, Han et al. 2012, Lee, Lee and Kim 2017, Gogolla et al. 2009). Additionally, low doses of the benzodiazepine clonazepam attenuated the impairments in sociability and cognitive function in autism rodent models (Han et al. 2012). However, the GABA_A_R subtypes involved in these effects are unknown. Here, we showed that the treatment with the α5-GABA_A_R positive allosteric modulator SH-053-2’F-R-CH3 attenuated deficits in social interaction and novel object recognition memory in males with no effect in female VPA rats, indicating sex differences for the effects of SH-053-2’F-R-CH3. This result is similar to previous studies describing sex differences for the anxiolytic-like effects of SH-053-2’F-R-CH3 (Piantadosi et al. 2016, Souza et al. 2022). This result does not seem to depend on differences in SH-053-2’F-R-CH3 bioavailability between males and females (Piantadosi et al. 2016), raising the possibility that the cause of sex differences in the behavioral effects induced by SH-053-2’F-R-CH3 may be associated with the effects, for example, of sex hormones. Thus, further studies should explore the effects of this compound at different stages of the estrous cycle.

α5-GABA_A_Rs are predominantly expressed on pyramidal cells in the hippocampus, and account for about 25% of hippocampal GABA_A_Rs (Olsen and Sieghart 2009, Hu et al. 2019). This restricted expression has attracted great interest in drug development in psychiatry (Jacob 2019). Preclinical studies indicate that positive allosteric modulators that act on the benzodiazepine site of α5-GABA_A_R, such as SH-053-2’F-R-CH3, produce anxiolytic-, antidepressant-, and antipsychotic-like effects (Prevot et al. 2019, Souza et al. 2022, Gill et al. 2011). Moreover, our findings agree with animal studies indicating that these compounds attenuate cognitive deficits (Prevot et al. 2019, Gill and Grace 2014).

In addition to the behavioral changes, we observed that *in utero* VPA exposure caused the adult animal, regardless of sex, to present an increased VTA DA neuron population activity. To our knowledge this is the first study investigating changes in the VTA DA system activity in the VPA autism model using in vivo electrophysiology. A hyperdopaminergic state has been extensively linked to psychotic disorders like schizophrenia. Notably, ASD share some alterations with schizophrenia, having previously been classified as an early form of schizophrenia with onset during childhood (KANNER 1949). Many young adults with schizophrenia also meet the DSM symptom criteria for ASD (Unenge Hallerbäck, Lugnegård and Gillberg 2012), and some ASD patients exhibit psychotic symptoms (Lugo Marín et al. 2018, Toal et al. 2009). Our findings are similar to the increases in VTA DA neuron population activity observed in animal models for schizophrenia, such as the one based on administering methylazoxymethanol acetate (MAM) to rats during gestation (Gill et al. 2011).

Despite attenuating behavioral changes, at least in males, SH-053-2’F-R-CH3 did not change the increased VTA DA neuron population activity in VPA rats. Contrary to our findings, SH-053-2’F-R-CH3 attenuated the hyperdopaminergic state in the MAM model (Gill et al. 2011). Other positive allosteric modulators of α5-GABA_A_Rs also reversed increases in VTA DA neuron population activity caused by stress and in an animal model for Alzheimer’s disease (McCoy et al. 2022, Eassa et al. 2023). In all these models, the VTA DA system overdrive is proposed to result from a ventral hippocampus hyperactivity due to a functional loss of local parvalbumin (PV)-containing GABAergic interneurons (Grace and Gomes 2019, McCoy et al. 2022, Eassa et al. 2023). Further studies are needed to evaluate whether an increased activity of the ventral hippocampus underly the increased VTA DA system activity in animals exposed to *in utero* VPA or it results from changes in the activity of other brain regions that also regulate the VTA DA system, such as the prefrontal cortex and thalamic nuclei (Zimmerman and Grace 2016, Patton, Bizup and Grace 2013).

Overall, our findings indicated SH-053-2’F-R-CH3 attenuates social and cognitive deficits in male animals exposed to VPA in utero without changing VTA DA system hyperactivity. Despite sex differences, these findings suggest that the GABAergic neurotransmission potentiation through positive allosteric modulators of α5-GABA_A_Rs may effectively attenuate core ASD symptoms.

## Acknowledgments

The authors thank Marco Antonio de Carvalho, Eleni Tamburus Gomes, and Eliane Aparecida Antunes Maciel for technical assistance. We also thank the Milwaukee Institute for Drug Discovery and the University of Wisconsin-Milwaukee’s Shimadzu Laboratory for Advanced and Applied Analytical Chemistry for help with spectroscopy and the National Science Foundation, Division of Chemistry [CHE-1625735].

## Contributors

AJS and FVG: conceptualization of the experiments, formal analysis, data analysis, and writing of the original draft. DS and JMC: synthesis of SH-053-2’F-R-CH3 and critically revised the manuscript. FVG: supervision. FSG: conceptualization of the experiments and critically revised the manuscript. All authors reviewed and approved the final version of the manuscript.

