## Supplementary figures for "An alpha 5-GABAa receptor positive allosteric modulator attenuates social and cognitive deficits without changing dopamine system hyperactivity in an animal model for autism"

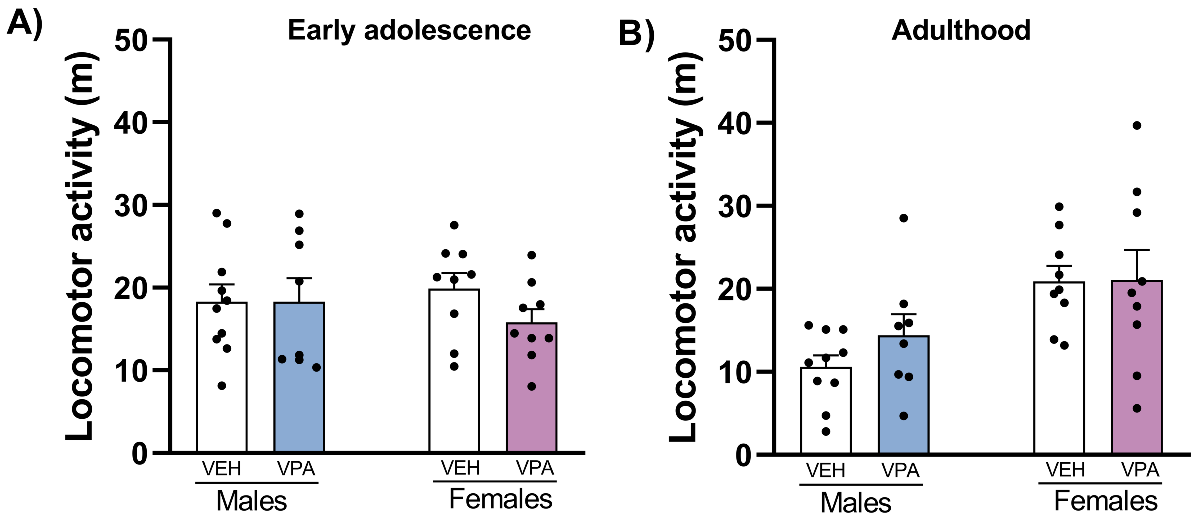


**Supplementary Figure 1** –*In utero* VPA exposure rats did not cause changes in the locomotor activity of males and females (A) during early adolescence (B) and adulthood (n=8-10/group).

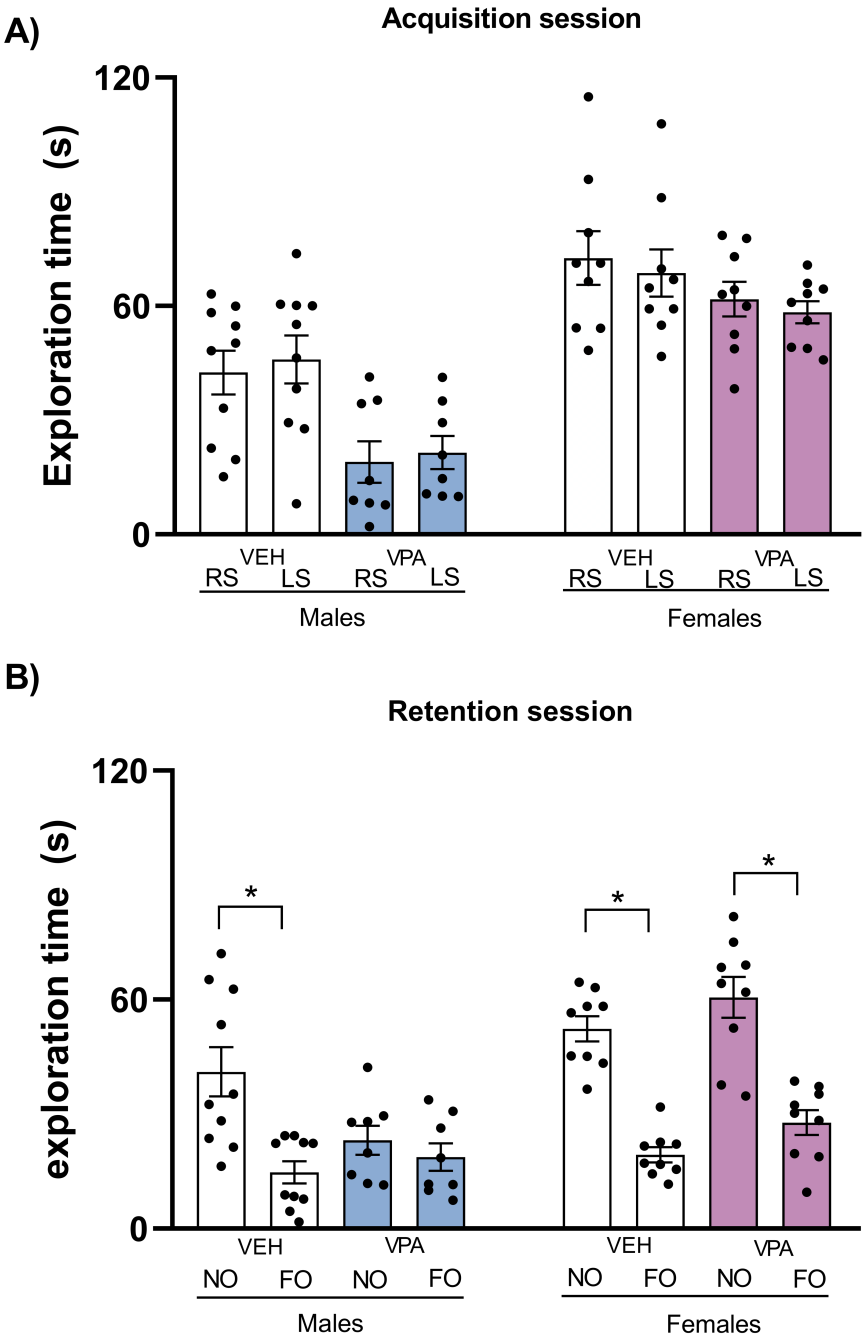


**Supplementary figure 2** – In the NOR test, (A) no difference between the exploration of the identical objects placed in the right (RS) or left side (LS) of the arena during the acquisition session was found indicating a lack of spatial preference. (B) In the retention session, a greater exploration of the novel object (NO) compared to the familiar object (FO) was observed for all groups, except for male adult rats exposed to VPA in utero (n=8-10/group). *p<0.05, Student t test.

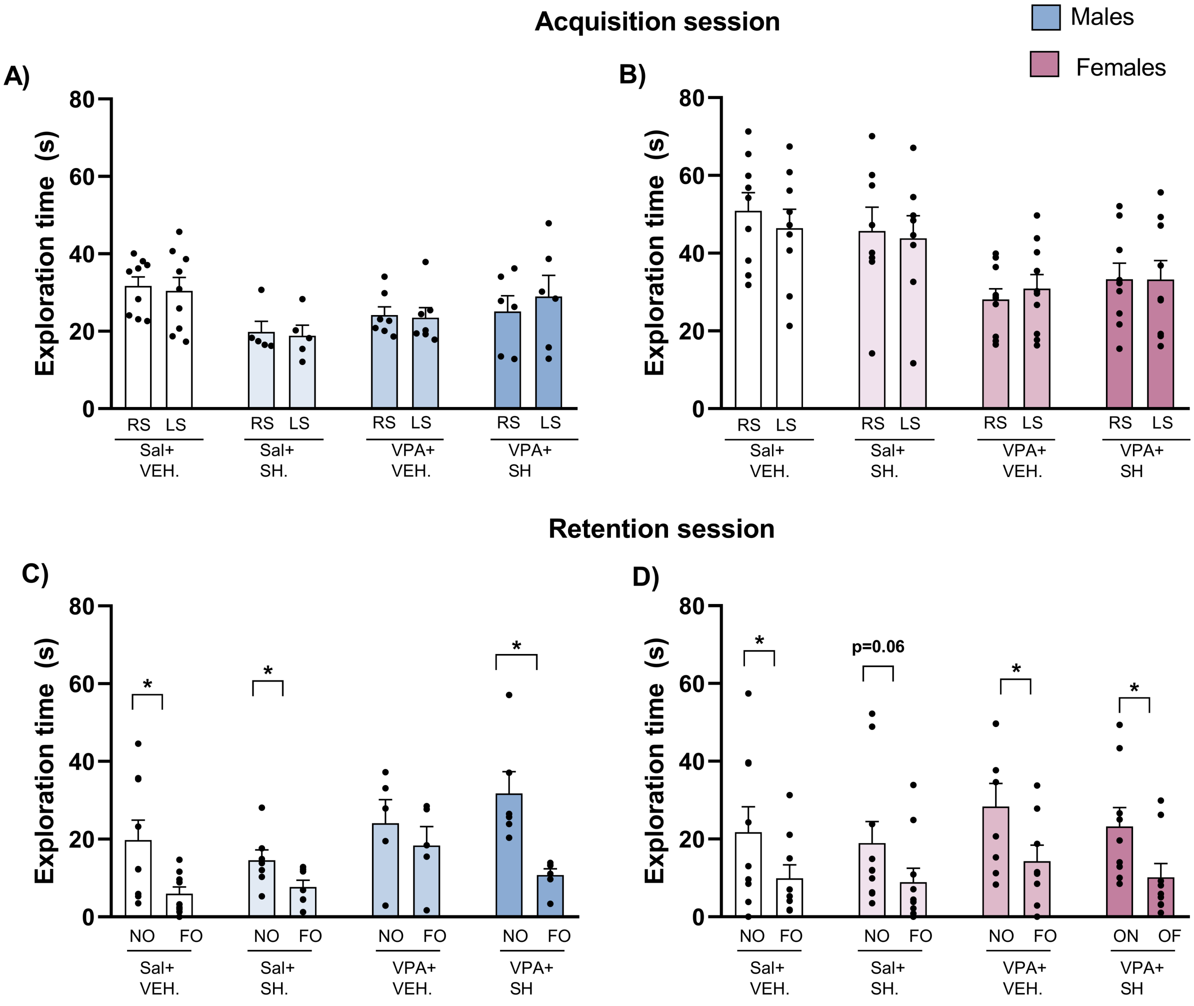


**Supplementary figure 3** – In the experiments with SH-053-2'F-R-CH3 (SH), no difference between the exploration of the identical objects placed in the right (RS) or left side (LS) of the arena during the acquisition session of the NOR test (A) for males (B) and females. C) In males, during the retention session, a greater exploration of the novel object (NO) compared to the familiar object (FO), was observed for all groups, except for those exposed to VPA. D) In females, a greater exploration of the novel object (NO), compare to the familiar object (FO), was observed for all groups (n=5-10/group) *p<0.05, Student t test.

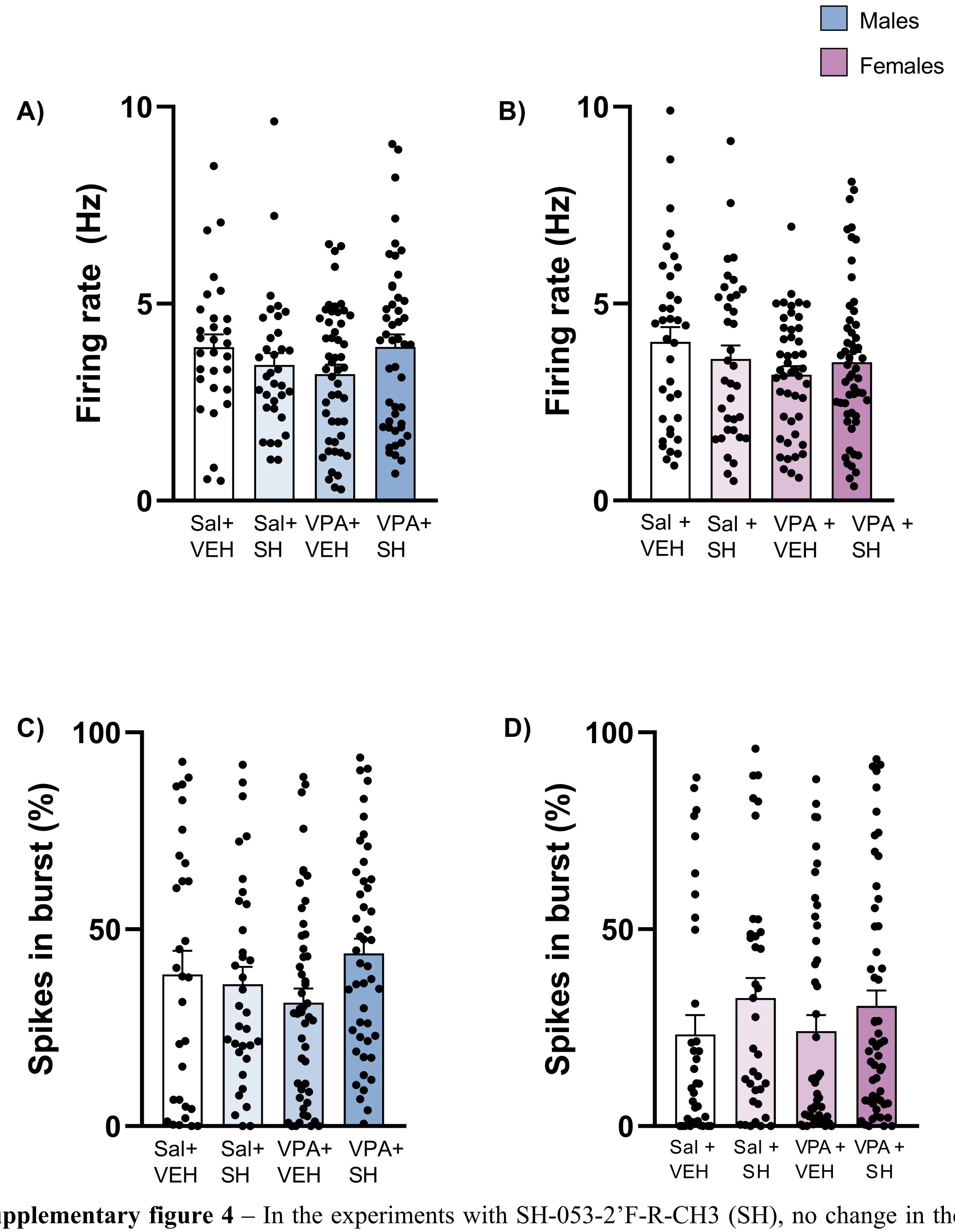


**Supplementary figure 4** – In the experiments with SH-053-2'F-R-CH3 (SH), no change in the (A-B) firing rate and (C-D) burst activity of VTA DA neurons (males: n= 40-54 cells/group; females: n=50-60 cells/group).
